# Gaze and perception covary with reading direction

**DOI:** 10.64898/2026.03.04.707660

**Authors:** Zahra Hussain, Riwa Safa, Rayan Kouzy, Julien Besle

## Abstract

Human observers prioritise the left visual field in perceptual judgements, a phenomenon widely attributed to cerebral asymmetries in vision. How stable are these mechanisms across differences in reading direction? We show that the classic left-side bias for faces is correlated with reading direction and tightly linked to gaze. Observers with varying proficiency in reading English (left-to-right) and Arabic (right-to-left) performed a chimeric face matching task (N = 271), with eye movements recorded in a subset (N = 115). The left-side bias scaled linearly with reading direction: strongest in English readers, reduced in bidirectional readers, and absent in Arabic readers. Reading direction also shifted gaze: English readers fixated further left on target faces and preferentially inspected faces in the left hemifield, and these gaze patterns predicted trial-by-trial perceptual judgements. These results suggest that cerebral asymmetries in face perception are flexible, coupled to gaze, and modified by reading direction.

---

By many accounts of looking and perceiving, the world is ordered from left to right. Gaze is directed first toward the left side of objects and scenes [1–3], spatial extent is overestimated in the left visual field [4], time and number progress from left to right in mental representations [5], and faces are judged primarily by the information in their left halves [6, 7]. Some of these leftward biases mirror asymmetric cerebral organization [7, 8]: the left visual field projects directly to the right hemisphere, which dominates in spatial processing and object recognition. Accordingly, leftward biases have been used to index cortical lateralisation [9, 10]. Yet experiential factors such as reading direction modify such biases and potentially their neural substrates [11–17]. Here, we ask whether reading direction guides perceptual bias and gaze in tandem during face perception, a process typically associated with right-hemispheric specialisation.

Robust leftward biases exist for faces. Perceived facial identity, age, emotion, attractiveness and gender are influenced more by what the left side of a face conveys, the side in the observer’s left visual field, than the right [7, 10, 18–20]. In a classic demonstration of this bias, chimeric faces are created from either the left or right half of a target face, and observers choose which chimera more closely matches the original (see Figure 1). The left chimera reliably is chosen more often than the right, even when the target is mirror reversed, indicating that the bias reflects visual field asymmetry rather than facial physiognomy. Consistent with the cerebral dominance hypothesis [7, 8, 10], the magnitude of the bias correlates with activation in right occipital-temporal face-selective cortex [fusiform face area; 21, 22]. Eye movements are likewise biased leftward [1, 20, 23], despite no evidence that the left side of the face is intrinsically more informative for recognition. Overall, this picture suggests a visual behaviour grounded in established neural asymmetries.

**Figure 1.**
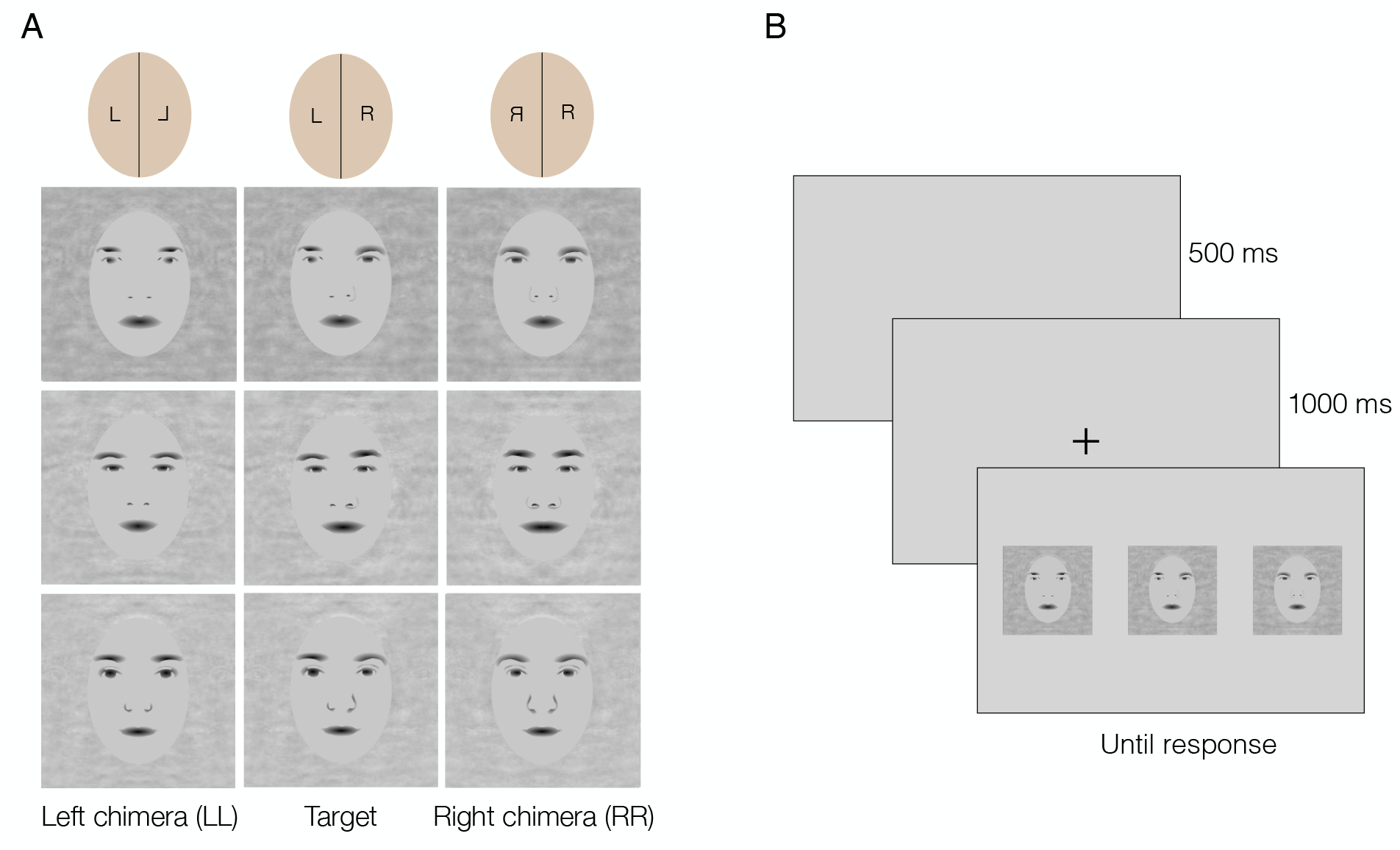
Examples of stimuli, shown here as caricatures to comply with bioRxiv policy. Real facial images were used for the experiment. (A) and schematic of the task (B).

Although the right-hemispheric dominance account is widely accepted, evidence suggests that reading direction contributes to leftward biases. In an early study, right-to-left Hebrew readers showed smaller and less reliable leftward biases in a chimeric face task than English readers [7], although this was interpreted as a universal bias at the time. Subsequent work has found the left-side bias for faces to be negligible or absent in readers of other right-to-left scripts, including Urdu and Arabic [11, 13, 14, 24]. Reading direction certainly affects biases in other spatial and perceptual judgements [15, 16, 25] and may fully account for directional biases in representations of time and number [26, 27].

Here, we examined the effect of reading direction on gaze and perceptual bias in a chimeric face resemblance task. First, we reproduced and extended the evidence for reading direction effects on perceptual bias in a large sample of left-to-right, right-to-left, and bidirectional readers. We then showed that reading direction also alters eye movements, and that gaze metrics predict perceptual judgements on a trial-by-trial basis.

## Materials and Methods

### Subjects

Subjects were 271 right-handed adults, of whom 115 participated in person with eye movements tracked, and 156 participated online, without eye-tracking. These sample sizes are commensurate with a previous study showing a small-to-medium effect of reading directionality on the left-side bias in a similar task [11, n = 131; Cohen’s f = 0.14, power > 0.9, calculated from Table 2 of this study]. A larger sample participated online to account for potential data loss from remote testing conditions, and to sample evenly across the continuum of reading directionality. Subjects were students and staff at the American University of Beirut (in person and online), and at McMaster University (online only). Arabic is the national language in Lebanon; however, English and French are widely spoken and taught in schools. Therefore, a large proportion of the Lebanese population is bilingual or multilingual. English- or French-only readers in this study largely comprised foreigners working or studying in Lebanon (having lived in English or French-speaking countries previously), and English-only speaking individuals in Lebanon. Arabic-only readers largely comprised custodial staff at the university. Bilingual or bidirectional readers largely comprised students and teaching staff at the university who could read both left-to-right (English or French) and right-to-left (Arabic) scripts. (The terms bilingual and bidirectional are used interchangeably in this work). All subjects completed a language proficiency questionnaire for Arabic and English (see Supplemental material). Subjects who participated in person also performed a reading task in their habitually-read language, either English or Arabic. Due to time constraints, bilingual readers did the task in any one language. Tables 1 & 2 provide subject details. The study was approved by the Institutional Review Board for Protection of Human Participants in Research and Research-Related Activities at the American University of Beirut. All subjects provided informed consent before they participated in the study. The final sample excluded two subjects with low reading proficiency in both Arabic and English, and eight subjects with poor eye-tracking data (281 subjects completed all tasks, and 10 subjects

**Table 1:** Participant information.

| **Location** | **Mean age (SD)** | **N. males** | **N. females** |
| --- | --- | --- | --- |
| In person | 32.5 (11.25) | 70 | 45 |
| Online | 19.6 (4.34) | 46 | 109 |

**Table 2:** Reading speed (mean words per second) of participants who performed the task in person, grouped by language (see text for criterion). Standard deviation and number of subjects in parentheses.

|  | English readers | Bidirectional readers | Arabic readers |
| --- | --- | --- | --- |
| English text | 3 (1.0, 22) | 3.1 (1.1, 24) | - |
| Arabic text | - | 2.3 (1.0, 17) | 2 (0.6, 52) |

### Language score

A language score for proficiency in English and Arabic was calculated for each subject based on self-reported reading, writing and speaking ability in both languages. The questions were five-point Likert questions developed in the lab (see Appendix). First, proficiency was scored for each subject in each language as a weighted average of responses to items 14-28 on the questionnaire, normalized between zero and one (no vs. high proficiency). Items related to reading ability were weighted more than other items. Figure 2A shows the distribution of proficiency scores as a scatterplot, with each point representing a subject’s score in the two languages. Monolingual readers are at the top left (Arabic) or bottom right (English) of the plot, and bilingual or bidirectional readers are at the top right. Absolute fluency is given by radial distance from the origin. Relative fluency in the two languages is given by angular position, with a polar angle of zero degrees corresponding to English-only, 90 degrees to Arabic-only, and varying degrees of bilingualism in between. Angular position was calculated for every subject, and rescaled to a score between −1 and 1, with zero denoting bidirectional readers. Henceforth, this score is referred to as the language score, and is used as a continuous predictor in the following analyses. For a subset of analyses or to simplify the illustrations, subjects were grouped as English readers (n = 95; < 35 deg), bidirectional readers (n = 121; 35 *−* 55 deg), and Arabic readers (n = 55; > 55 deg). Two subjects not sufficiently fluent in either language (shown by triangles in Figure 2A; absolute fluency < 0.75) were excluded from the analyses.

**Figure 2.**
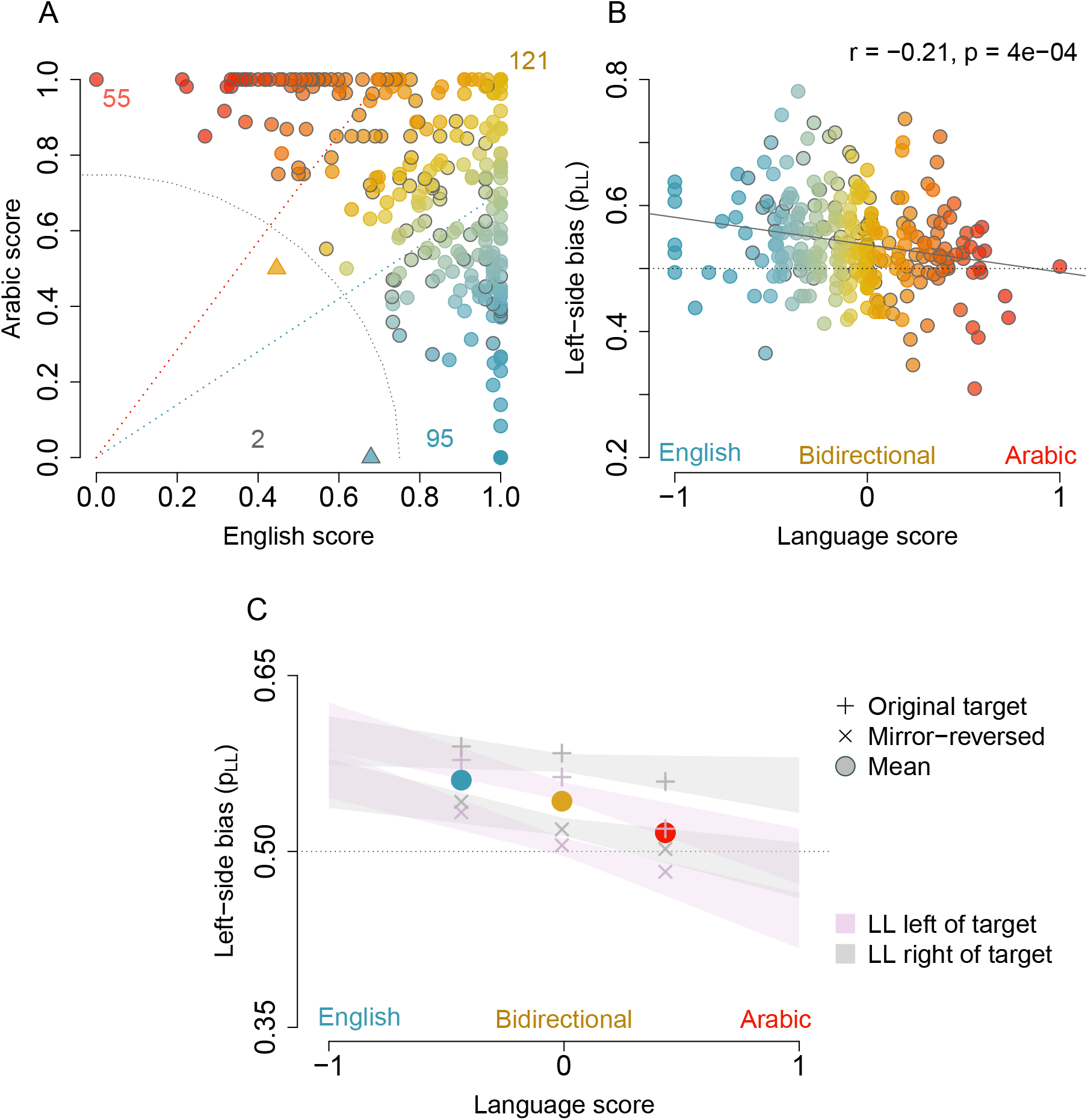
A) Scatterplot of language scores for the full sample of 271 subjects. Symbol colour varies with relative proficiency in two languages (red: monolingual Arabic; blue: monolingual English; yellowish: bilingual). Dashed diagonal lines show criteria for group assignment (polar angles of 35 deg and 55 deg). Dotted grey line indicates fluency criterion for inclusion in the study (r = 0.75 in these coordinates). Numbers indicate N per group. Symbols outlined in grey: eye-tracking subjects. B) The left-side bias averaged over stimulus conditions against language score for the full sample. Solid grey line: best linear fit. C) The left-side bias in each stimulus condition for the three groups. Shaded regions show the standard error of the predicted effects of language score, target type (original vs. mirror-reversed), and LL location (left vs. right visual field). Plus vs. cross symbols: Original vs. mirror-reversed target. Rose vs. grey shading: LL in left vs. right visual field.

### Reading speed

Subjects who performed the tasks in person did a reading speed task prior to the main experiment. This was done to confirm reading ability, and to evaluate potential differences in reading ability across groups. In the reading speed task, subjects were shown a passage of English or Arabic text on the screen (English: 242 words; Arabic: 158 words), with instructions to press a spacebar once they had read the passage. This was followed by four questions that probed understanding of the material. Reading speed was measured as the number of words in the initial passage divided by the time taken to press the spacebar in seconds. Table 2 gives the mean reading speeds for subjects grouped by language. Reading speeds were higher on average for English text than Arabic text, but equivalent across groups within each language. A t test confirmed that reading speed for English text did not differ significantly between English and bidirectional readers (*t*(44) = 0.23, *p* = 0.83), and that reading speed for Arabic text did not differ significantly between Arabic and bidirectional readers (*t*(20.97) = 1.07, *p* = 0.29). Note that bidirectional readers did the reading task in only one language, so the above comparisons were across different subjects.

### Apparatus and Stimuli

In person, the task was performed on a Tobii T120 eyetracker at a sampling rate of 120 Hz, on a computer screen set to a resolution of 1280 *×* 1024 pixels. Online, the task was performed on subjects’ own computers on Pavlovia, a platform for psychology experiments (https://pavlovia.org/). Viewing distance was set to 57 cm for the eyetracking subjects, and assumed to be similar, but was not monitored for the online subjects.

The stimuli were 40 greyscale male and female Caucasian faces with neutral facial expressions, cropped to show only internal features within an oval subtending 190:140 pixels, and equated in amplitude spectra. The faces were presented in a square frame (256 *×* 256 pixels), subtending approximately 5 x 5 degrees of visual angle from a viewing distance of 57 cm. The stimuli were derived from a stimulus set created by colleagues for a former study [28]. For each target face, a left and right chimeric version was created by splitting the face down its vertical midline and recombining the left and right halves with their mirror-reversed versions (Figure 1). A mirror-reversed version of the target was also created. For both original and mirror-reversed targets, the left chimera (LL) was the face comprised of the half in the observer’s left visual field, and the right chimera (RR), was the face comprised of the half in the observer’s right visual field. That is, LL for the original target served as RR for the mirror-reversed target. The full stimulus set comprised 160 faces: four versions of each of the 40 faces (original, LL, RR, mirror-reversed). For a few faces, the LL and RR halves were adjusted to be one or two pixels less or greater in width than the exact midpoint of the face to avoid artefacts.

### Procedure

Subjects provided informed consent and completed a language and handedness questionnaire before performing the experimental task (Appendix). Subjects who participated in person were then seated in a soundproof booth in front of the eye-tracker. Subjects who participated online were provided with the link to the experiment and instructions on the conditions in which to perform the task (on a laptop or desktop computer using Google Chrome). For the in-person subjects, the session began with an eye-tracking calibration procedure, followed by the experimental task. The task parameters were the same for all subjects. Subjects who provided incomplete responses to the questionnaire or who did not complete all trials, were excluded from the study.

Each trial in the experimental task began with a blank screen (500 ms), followed by a fixation cross in the centre of the screen (1 s). The fixation cross was followed by a set of three faces that remained on the screen until the subject responded. The face at the centre of the screen was always the target face, and the type of target (original or mirror-reversed) was randomised across trials. The chimeric choice faces (LL, RR) were positioned 6 degrees to the left and right of the target, with LL and RR appearing equally often and randomly on either side of the target. Subjects used a keypress to report which chimeric option more closely resembled the target. Subjects were unaware of the chimeric face manipulation, and were free to move their eyes whilst making the judgement. In-person subjects performed two trials per condition for a total of 320 trials (40 faces *×* 2 target types (original, mirror-reversed) *×* 2 visual field positions for each chimeric face (left, right) *×* 2 iterations). Online subjects did one trial per condition or 160 trials in all.

### Analysis

The data were analyzed using R software for statistical computing [29].

### Perceptual bias

The left-side bias, *pLL*, was computed as the proportion of trials on which subjects chose the left chimera (LL) as a better match to the target. By this measure, 50% indicates no bias, and values significantly above and below 50% indicate left- and right-sided biases. For trial-wise analyses, the binary outcomes (LL chosen = 1, RR chosen = 0) were analysed.

### Eye-tracking

The eye-tracking analyses focused on the period between the onset and offset of the face stimuli, which depended on subjects’ response time. Trials were excluded if more than 10% of eye-tracking data samples were missing during this time window (e.g., due to blinking or poor calibration), and subjects were excluded if more than half of the 320 trials were removed due to missing samples. Six subjects were excluded on this basis, and a further two subjects were excluded because their gaze was consistently centred outside the face stimuli. Hence, 115 subjects were included in the eye-tracking analyses.

For each subject, saccades and fixations were extracted on each trial using the following procedure: first, the instantaneous gaze position was temporally smoothed using a 5-sample (41.67 ms) window. Next, saccades were identified as groups of more than 2 consecutive samples with velocities (changes in gaze position) over 840 pixels/s, and fixations were identified as groups of consecutive samples between saccades. Summary fixation and saccade measures were then computed for the three faces (left, target, and right face) for each subject on each trial. The average fixation coordinates on each face were calculated by first averaging the coordinate values across samples within each fixation, and then across fixations within each trial. Coordinates were averaged between the left and right eyes at each sample, and expressed as the distance from the centre of the face in degrees of visual angle. We also calculated the earliest saccade number at which gaze was directed to the choice faces after inspection of the target, and the number of fixations within each face.

### Models

Statistical tests included linear mixed-effects (lmer) and generalized linear mixed-effects (glmer) models implemented in the lme4 and lmerTest packages [30, 31]). These approaches are appropriate for designs combining continuous and categorical predictors.

Perceptual bias was analysed on subject-level aggregated data using lmer, with *pLL* as the dependent measure. The model included language score as a continuous fixed predictor, target version (original vs. mirror-reversed) and LL location (left vs. right hemifield) as categorical fixed effects, and random intercepts for subject. These analyses were conducted on the full sample comprising 271 subjects.

Gaze attributes also were analysed on aggregated data using lmer, with fixation coordinates (*x* or *y*), saccade order, or number of fixations as the dependent measures. Fixed effects included language score and face (left vs. centre. vs. right, or left vs. right depending on the analysis), and random intercepts were included for subject. Stimulus variables (target version and LL location) were dropped from these models after initial analyses showed that they did not account for significant variance. These analyses were conducted on the subset of 115 subjects who performed the task in person.

Perception-gaze associations were analysed at the trial-level using (glmer) with a logit link, where the dependent measure was LL success (0/1). Success was defined as the selection of the LL chimera as a match to the target (1), and failure as the selection of the RR chimera (0). The significance of fixed effects was evaluated with Type III Wald *χ*^2^ tests. Trial-level models were used to gauge the dynamic effects of gaze attributes on perceptual choice.

For the lmer models, partial eta-squared 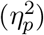 is reported as a measure of effect size. For glmer models, effect sizes are reported as odds ratios with 95% confidence intervals.

## Results

### Perceptual bias

Figure 2B shows the left-side bias, *pLL*, against language score for each participant, on average over the four stimulus conditions (target version: original vs. mirror-reversed target; LL location: left vs. right hemifield). A simple linear model confirmed that *pLL* was correlated with reading directionality, positive for English readers and negligible or absent for Arabic readers (*r*(269) = *−*0.21, *p* = 0.0004). A two-tailed t-test confirmed that *pLL* was significantly above chance or 50% for the sample overall (*t*(270) = 9.36, *p* < 0.0001;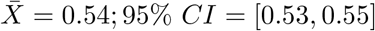).

Figure 2C shows the left-side bias in the four stimulus conditions. The shaded regions represent the predicted effects of language score on *pLL* from the glme logit model. The symbols show group means of *pLL* in each stimulus condition at the mean language score for each group (as identified in Figure 2A). The figure confirms the inverse relationship between left-side bias and Arabic reading score shown in Figure 2B, and reveals an unexpected effect of target type: the bias was stronger for original targets (plus symbols) than mirror-reversed targets (crosses) for all groups. Additionally, the bias was greater when the LL choice appeared to the right of the target. These effects were confirmed by significant main effects of language score (*F* (1, 269) = 12.83, *p* = 0.0004, 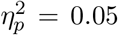), target version (*F* (1, 807) = 51.28, *p* < 0.00001, 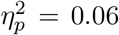), and hemifield (*F* (1, 807) = 8.50, *p* = 0.004, 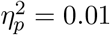). None of the interaction effects were significant (*F* < 2.5, *p >* 0.12). The fixed effects explained about 6.2% of the variance in bias (marginal *R*^2^), while the full model including random effects explained about 17.0% of the variance (conditional *R*^2^). In other words, there was substantial individual variation in bias.

In summary, the left-side bias for faces was reduced in Arabic readers compared to English readers, smaller for mirror-reversed targets than original targets, and larger when the LL chimera appeared in the right visual field.

### Gaze patterns

#### Fixation coordinates

Figure 3 shows the spread of fixation coordinates on the face stimuli across all trials for three representative observers in each language group. Ninety-five percent confidence intervals for each subject’s 2D fixation coordinates are shown as ellipses. Fixations were both idiosyncratic and strikingly consistent across trials, with ellipse area around half a degree of visual angle for the target face, and slightly larger for the choice faces. Box plots at the bottom of the figure display the distribution of ellipse area for the full sample, indicating that fixation consistency (or spread) was roughly comparable across the three language groups, with English readers slightly more consistent than the others.

**Figure 3.**
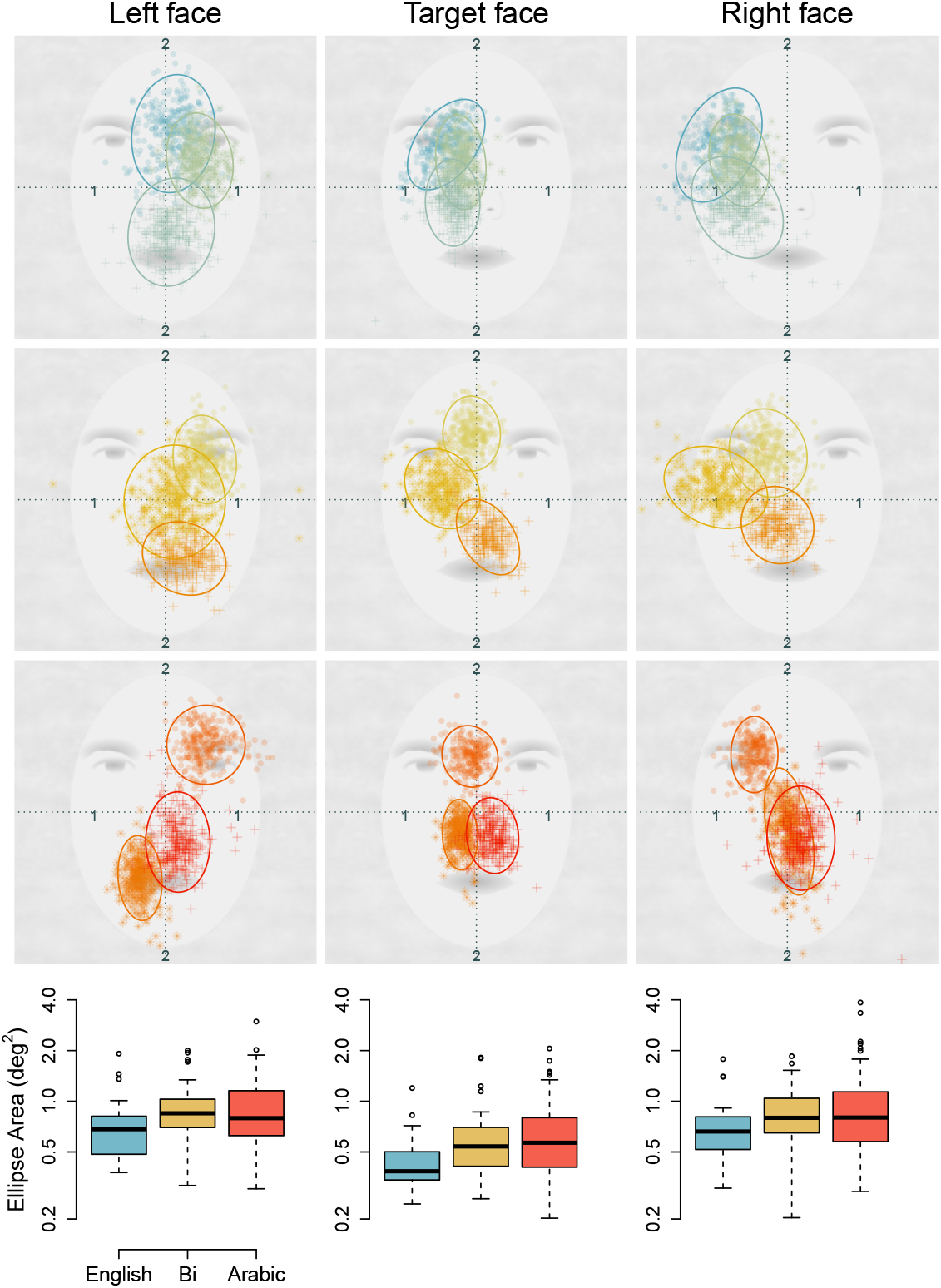
Fixation coordinates within the three faces for three representative subjects in each language group (top: English, middle: Bidirectional, bottom: Arabic), superimposed on caricature face stimuli. Numbers show coordinates in degrees of visual angle. Symbols show average fixation x- and y location across fixation samples within a face per subject per trial. The number of trials (or fixations) per subject per face ranges from 246 to 318. Ellipses show 95% confidence intervals for each subject’s 2D fixation coordinates across all trials. Bottom row: Box plots of ellipse area for each face across all subjects in each group.

Biases in fixation location are shown in Figure 4. Figure 4A shows average fixation positions for the three groups in each of the three faces during the choice period. These coordinates are averaged over samples, fixations (median = 2-3 fixations per face), trials, stimulus conditions (target type and LL location), and subjects. On average across faces, English readers’ fixations were farther leftward compared to the other groups, and Arabic readers fixated closer to the centre of the face. A screen-centre bias is evident in all groups: fixations on the left face were rightward of midline, and fixations on the right face were leftward. Hence, group, or language-related differences are clearest in the target face. Vertically, fixations were above the midpoint of the image (i.e., closer to the eyes) in most conditions.

**Figure 4.**
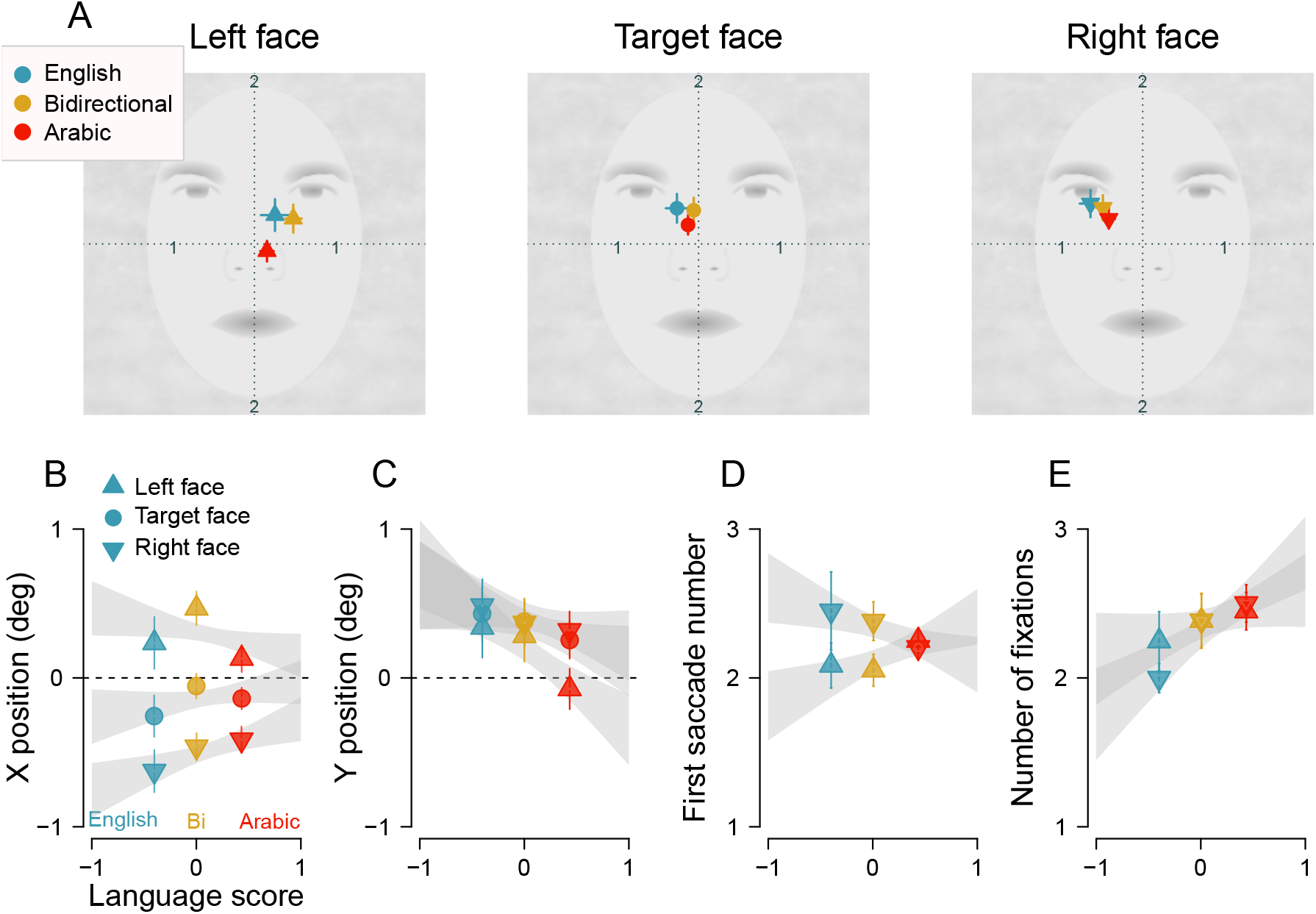
A. Average gaze positions on the three caricature face stimuli during the choice period for the three language groups. Sample sizes were 22 (English), 41 (Bidirectional), and 52 (Arabic). Dotted lines indicate face-centered coordinates and numbers in black mark degrees of visual angle (not shown in the experiment). B-E. Predicted effects of language score on gaze x- and y-coordinates (B & C), first fixation number for the left vs. right choice image (D), and number of fixations (E; see text for model details). Symbols show group means, at the mean language score for each group. Error bars and shaded regions indicate standard error of the mean.

The fixation position biases are confirmed in Figures 4B & C, which show the predicted effects of language score on fixation x- and y-coordinates for the three faces, based on lme models fit to the coordinate data aggregated over target version and hemifield. For fixation x-coordinates (Figure 4B), there was a significant main effect of face (*F* (2, 226) = 139.84, *p* < 0.0001, 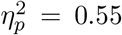) reflecting the horizontal screen-centre bias described above. There was a significant interaction between language score and face (*F* (2, 226) = 4.54, *p* = 0.012, 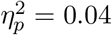), supporting the observation that English reading was associated with larger horizontal biases on the three faces (i.e., leftward on the target and right face, and rightward on the left face), compared to Arabic reading. The main effect of language was not significant (*F* (1, 113) = 0.23, *p* = 0.63). Marginal *R*^2^ and conditional *R*^2^ were 24% and 71.4%.

For fixation y-coordinates (Figure 4C), there was a significant main effect of face (*F* (2, 226) = 38.96, *p* < 0.0001, 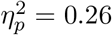), confirming that fixations were higher on the target and right face than on the left face. There was a significant language *×* face interaction (*F* (2, 226) = 15.83, *p* < 0.0001, 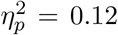), confirming that English reading was associated with higher fixations than Arabic reading, especially for the left face. The main effect of language was not significant (*F* (1, 113) = 1.86, *p* = 0.18). Marginal *R*^2^ and conditional *R*^2^ were 3.2% and 96%. Hence, there was substantial individual variation in both horizontal and vertical fixation bias.

#### Saccade sequence: left vs. right face

Figure 4D plots the saccade number at which participants first looked at the left and right faces, as a function of language score. The figure shows that English and bidirectional readers tended to look at the left face before the right, whereas Arabic readers did not show this preference. Averaged over groups, the left face was inspected first. Consistent with these patterns, there was a significant main effect of face (left vs. right; *F* (1, 113) = 5.57, *p* = 0.019,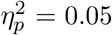), and significant interaction between language score and face (*F* (1, 113) = 4.417, *p* = 0.04, 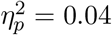). The main effect of language score was not significant (*F* (1, 113) = 0.02, *p* = 0.88). Marginal *R*^2^ and conditional *R*^2^ were 3% and 8%.

#### Number of fixations: left vs. right face

Figure 4E plots the number of fixations on each face as a function of language score. English readers made more fixations on the left face than the right, whereas bidirectional and Arabic readers did not show this asymmetry. This effect was confirmed by a significant interaction between language score and face (*F* (1, 113) = 5.41, *p* = 0.02, 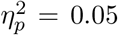). The main effects of language score and face were not significant (*F* < 2.4, *p >* 0.14). Marginal *R*^2^ and conditional *R*^2^ were 2% and 87%.

In summary, larger leftward biases were found in English readers compared to Arabic readers on several gaze measures, including fixation location on the target face, first saccade toward the left or right choice face, and number of fixations on the left vs. right choice face.

#### Gaze effects on perception

We examined trial-level gaze dynamics by testing whether the left-side bias on individual trials was linked to fixation patterns on the target and choice faces. Choice faces were coded by chimera type (LL vs. RR) rather than screen location. glmer logit models were used, with the binary outcome on each trial as the dependent measure. Separate models tested the effects of (i) fixation position on the target face, (ii) frequency of looks to the choice faces (LL vs. RR), and (iii) the order of fixations on the choice faces, with language score included as a continuous predictor. Random intercepts for subjects were included in all models. Each model tested whether these gaze measures predicted perceptual choice on a trial-by-trial basis.

Figure 5A shows the predicted effect of fixation x-position on the target face. A clear linear relationship was found: leftward fixations were associated with a stronger left-side bias across all subjects, regardless of reading direction (small grey symbols represent the effect of x-coordinate averaged over language score). These effects were confirmed by a significant main effect of x-coordinate (*χ*^2^(1) = 33.7, *p* < 0.00001), and a significant main effect of language score (*χ*^2^(1) = 9.61, *p* = 0.0019), consistent with earlier results. Larger x-coordinate values were associated with lower odds of choosing the LL chimera (*OR* = 0.79, 95% CI [0.73, 0.86]), as were increases in language score (*OR* = 0.75, 95% CI [0.62, 0.90]). The interaction between x-coordinate and language was not significant (*χ*^2^(1) = 0.006, *p* = 0.94). Thus, leftward fixations on the target face predicted a stronger left-side bias across all readers.

**Figure 5.**
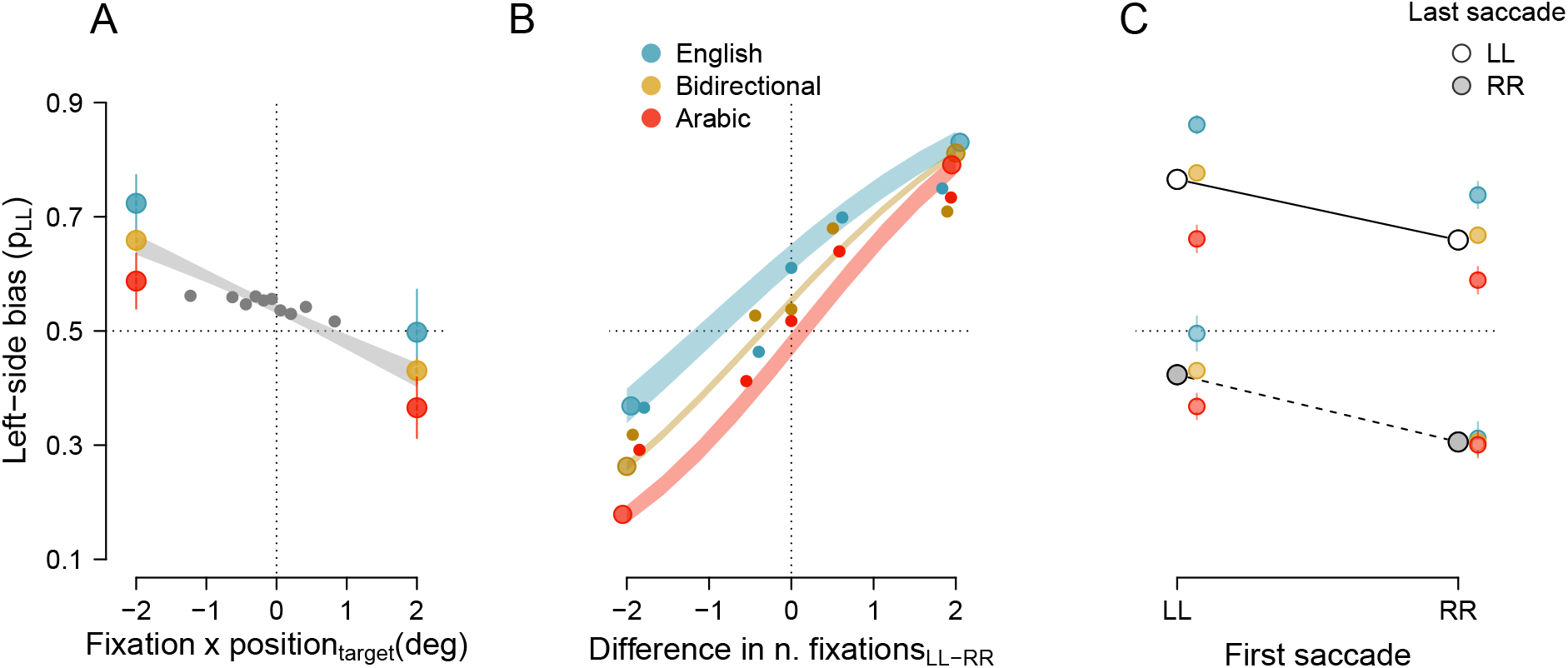
A. The left-side bias as a function of fixation x position on the target face. Shaded regions and error bars show standard error of the mean of predicted effects. Coloured symbols show the predicted bias for each language group at the two extreme fixation positions (reflecting the main effect of language score and no interaction of language with fixation position; see text). Smaller grey symbols show the mean bias across language score at ten quantiles of fixation position. B. The left-side bias as a function of the difference in number of fixations between the LL and RR choice faces, shown separately for the three language groups (reflecting the interaction of language score with difference in number of fixations; see text). Shaded regions show standard error of predicted bias. Large symbols show predicted effects; smaller symbols show bias computed at five quantiles of fix-difference. C. The left-side bias as a function of fixation order (first vs. last) on the LL and RR stimuli. Abscissa shows bias separately for first fixations directed to the LL or RR face. Different symbols (open vs. grey) show bias separately for the last fixation stimulus (LL vs. RR).

Figure 5B shows the predicted effect on the left-side bias of the difference in the number of fixations on the LL and RR choice faces (irrespective of which visual field they were in). For each subject, on each trial, we computed the difference between the number of fixations on the LL versus the RR face (fix-difference), and the z-scaled sum of those fixations (fix-sum; scaling improved model convergence). These orthogonal measures were used as fixed predictors, along with language score, of trial-wise choices on the behavioural task. The model revealed significant main effects of fix-difference (*χ*^2^(1) = 2311.88, *p* < 0.00001), fix-sum (*χ*^2^(1) = 7.76, *p* = 0.005), language (*χ*^2^(1) = 12.57, *p* = 0.0003), and significant interactions between fix-difference and fix-sum (*χ*^2^(1) = 621.56, *p* < 0.00001), and between fix-difference and language (*χ*^2^(1) = 6.86, *p* = 0.008). Figure 5B shows the fix-difference *×* language interaction and main effects, with language plotted at three levels (groups). There was a monotonic increase in the left-side bias as the number of fixations on the LL face exceeded the number of fixations on the RR face (fix-diff *OR* = 1.86, 95% CI [1.82, 1.91]), and this effect was smaller for English readers (i.e., the slope of the relationship was shallower; fix-diff *×* language *OR* = 1.09, 95% CI [1.02, 1.17]). In other words, the left-side bias depended on the frequency of looking at the choice faces. The fix-difference *×* fix-sum interaction reflected the finding that a difference in the number of fixations on LL versus RR faces had a greater impact when there were few total fixations in a trial compared to when there were many fixations (data not shown). In other words, the slope of the bias vs. fix-difference curve was steeper when there were few total fixations in a trial (fix-diff *×* fix-sum *OR* = 0.82, 95% CI [0.81, 0.84]).

Figure 5C shows the predicted effect on the left-side bias of the order of looking at the LL versus the RR face. For each subject, on each trial, binary variables were computed to identify whether the first saccade and the last saccade was made to the LL face. These variables were used as predictors along with language score, of trial-wise choices on the behavioural task. Figure 5C shows that the left-side bias was larger both when the first saccade was to the LL face, and when the last saccade was to the LL face, with the last saccade exerting a larger effect. There were significant main effects of first saccade (*χ*^2^(1) = 202.69, *p* < 0.00001, *OR* = 1.71, 95% CI [1.59, 1.84]) and last saccade (*χ*^2^(1) = 1610.44, *p* < 0.00001, *OR* = 4.551, 95% CI [4.22, 4.90]), and significant interactions between language score and first saccade (*χ*^2^(1) = 5.23, *p* = 0.02, *OR* = 0.79, 95% CI [0.65, 0.96]), and language score and last saccade (*χ*^2^(1) = 9.09, *p* =, *OR* = 0.73, 95% CI [0.60, 0.90]). These interactions reflected the fact that the language score effect was absent when neither the first nor the last saccade was made to the LL face (i.e., when the RR face drew more attention). In those circumstances, there was a right-side bias instead of a left-side bias for all groups. Hence, the timing or order of inspection of the choice faces predicted the left-side bias. The main effects of language and the other interactions were not significant (*p >* 0.8).

In summary, these analyses show robust associations between eye movements toward the target face and choice faces, and perceptual decisions on the chimeric face task. Specifically, the likelihood of a left-side bias increased with leftward fixations on the target face, when there were relatively more fixations on the LL choice face than on the RR face, and if the first and final saccades on a trial were directed to the LL face instead of the RR face.

## Discussion

Wolff [6] first reported that individuals failed to recognise their right-hand selves when viewing chimeric photographs of their own faces, whereas strangers failed to identify left-hand versions of those individuals. Many studies have since confirmed this left-side bias for faces and attributed it to specialised, possibly innate mechanisms in the right cerebral hemisphere [7, 10, 18–20, 32]. Reading experience can recalibrate such mechanisms [11, 13, 14, 24, 33], and we have shown that it concurrently reallocates gaze in step with perceptual changes. These effects resemble forms of experience-dependent plasticity [e.g., perceptual learning; 34, 35], particularly acquired expertise that generalises beyond the trained domain and alters brain organisation [36–42].

The perceptual bias for English readers (5-10% above chance) was similar in magnitude to previous studies, and it was absent for Arabic readers. That Arabic readers did not show an opposite, right-side bias, is consistent with reports of left-side biases in preliterate children and infants [43–45], suggesting the leftward bias is at least partly innate. Unexpectedly, the bias was stronger in all subjects for the original target faces compared to mirror-reversed targets. Mirror-reversed faces are typically used to show that the bias is unrelated to asymmetries in facial physiognomy, but in our data, facial or image-specific differences may have contributed. Well-known asymmetries in facial expression and musculature could direct attention toward one side of the face [46, 47]. However, facial emotion is generally expressed more strongly on the left physical side (observer’s right visual field), opposite to the observed bias, due to right-hemispheric dominance in emotion processing combined with contralateral facial control [47, 48]. One possible interpretation is that the visual field bias emerges to counterbalance these asymmetries in emotional expression between the two facial halves.

In oculomotor behaviour, English readers oriented further leftward overall, fixating to the left on the target face, and inspecting the left choice face more frequently or earlier than the right. This pattern matches previous reports of leftward gaze biases for faces [1, 23]. The effect of reading direction on horizontal fixation emerged despite highly idiosyncratic looking patterns for faces which have been reported previously [49], and which were evident here (Figure 3; the random effect of subject accounted for substantial variance in fixation position). Arabic readers showed no directional biases on any measures, again supporting the view that reading experience modifies an initially leftward bias. While the absence of leftward biases in Arabic readers could be interpreted as a general cultural difference [e.g., 50], our results suggest it is more specifically linked to reading experience. For other factors to account for these findings, they would need to correlate with language proficiency (or correlate with the perceptual bias as strongly as language score does). A potential caveat is that the Arabic-only readers in this sample were older than other readers, but the left-side bias is uncorrelated with age [51], and data from the bidirectional-readers suggest that age was not a confound.

The covariation between gaze parameters and perceptual choice in this task suggests that judgements of facial appearance were based on the specific facial regions sampled during eye movements. The left-side bias was related linearly to the horizontal coordinates of fixations on the target face, dropping to zero as fixations shifted rightward. This provides the clearest evidence, as far as we know, of perception so tightly linked to gaze location on a face stimulus. Moreover, the bias was associated with both a greater number of fixations on the left chimera and earlier saccades toward it. Previous studies have reported longer or more frequent fixations toward the left side of a chimeric face, on average across left-bias trials [20, 51]. These covariations between oculomotor behaviour and perception are consistent with with evidence that collicular mechanisms contribute to perceptual decision making [52–54], and that oculomotor signals are plastic [55]. For faces, preexisting bias may guide gaze toward the preferred visual field, while task-dependent gaze dynamics and stimulus attributes reciprocally influence perceptual choice.

Precisely what neural changes could reading direction induce, and how might they be coordinated across subcortical and cortical mechanisms? Studies of word recognition offer some clues. Roman-script readers versus Hebrew or Arabic readers show opposite asymmetric perceptual spans, viewing position effects, and preferred fixation landing sites within words [left vs. right of center; 56, 57], and, less consistently, opposite visual field advantages in word recognition [right vs. left visual field; 35, 58]. These differences have been interpreted as reading-induced modifications in early visual areas corresponding to retinally-specific pattern learning [35], as well as changes further upstream, in lateralised regions subserving printed word recognition, such as the visual word form area [VWFA; 59]. It has been proposed that cortical asymmetries for reading and face recognition co-develop [60–63], and such development may depend on script properties. We currently are examining the lateralisation of neural representations for faces and words in Arabic compared to English readers.

## Acknowledgments

This work was funded by a University Research Board grant to Zahra Hussain at the American University of Beirut.

## Supplemental Material

**Table 3:** Demographic, language and handedness questionnaire.

| Question | Options / subitems |
| --- | --- |
| 1. Sex |  |
| 2. Age |  |
| 3. Handedness | left vs. right |
| 4. Vision | normal vs. corrected vs. impaired |
| 5. Hearing | normal vs. corrected vs. impaired |
| 6. Have you ever had any language or learning disability? | If yes, please explain |
| 7. Current occupation |  |
| 8. What is your highest level of education? | less than high school; high school; professional training; undergraduate education; master's; PhD/MD/JD; Other |
| 9. What was the language of instruction at your highest level of education? | English; Arabic; French; Other |
| 10. Have you ever lived in a country where Arabic is NOT the dominant communicating language? | If yes, for how long? |
| 11. Have you ever lived in a country where Arabic is the dominant communicating language? | If yes, for how long? |
| 12. List three languages you know in order of fluency |  |
| 13. On a scale of 0 (not proficient) to 4 (highly proficient) rate your proficiency in your fluent language with a script written from **left to right** (e.g., English, French, Hindi): | *speaking* |
| 14. | *writing* |
| 15. | *reading* |
| 16. | *understanding* |
| 17. On a scale of 0 (not proficient) to 4 (highly proficient) rate your proficiency in your fluent language with a script written from **right to left** (e.g., Arabic, Hebrew, Urdu): | *speaking* |
| 18. | *writing* |
| 19. | *reading* |
| 20. | *understanding* |
| 21. On a scale of 0 to 4, how much of the following do you do in a language with a script written from **left to right** (e.g., English, French, Hindi): | *speaking* |
| 22. | *listening* |
| 23. | *writing* |
| 24. | *reading* |
| 25. On a scale of 0 to 4, how much of the following do you do in a language with a script written from **right to left** (e.g., Arabic, Hebrew, Urdu): | *speaking* |
| 26. | *listening* |
| 27. | *writing* |
| 28. | *reading* |
| 29. For English or another language with a script written from **left to right**, what kind of material do you read, and how many hours on each type of material (choose all that apply) | books; newspapers; encyclopaediae; religious texts; magazines; academic papers; academic books; online articles/blog posts; other |
| 30. Same as above for Arabic or other **right to left** script |  |
| 31. On a scale of 1 to 5, which hand do you prefer to use for: | *writing* |
| 32. | *throwing* |
| 33. | *using a toothbrush* |
| 34. | *using a spoon* |

